# Genotypic variation in morphological source and sink traits affects the response of rice photosynthesis and growth to elevated atmospheric CO_2_

**DOI:** 10.1101/694307

**Authors:** Denis Fabre, Michael Dingkuhn, Xinyou Yin, Anne Clément-Vidal, Sandrine Roques, Armelle Soutiras, Delphine Luquet

**Author notes:** Email address of each authors. **Corresponding author:** Denis Fabre, CIRAD, UMR AGAP, F-34398 Montpellier, France.

## Abstract

This study aimed to understand the response of photosynthesis and growth to e-CO_2_ conditions (800 vs. 400 μmol mol^-1^) of rice genotypes differing in source-sink relationships. A proxy trait called local C source-sink ratio was defined as the ratio of flag leaf area over the number of spikelets on the corresponding panicle, and five genotypes differing in this ratio were grown in a controlled greenhouse. Differential CO_2_ resources were applied either during the two weeks following heading (EXP1) or during the whole growth cycle (EXP2). Under e-CO_2_, low source-sink ratio cultivars (LSS) had greater gains in photosynthesis, and they accumulated less nonstructural carbohydrate in the flag leaf than high source-sink ratio cultivars (HSS). In EXP2, grain yield and biomass gain was also greater in LSS probably caused by their strong sink. Photosynthetic capacity response to e-CO_2_ was negatively correlated across genotypes with local C source-sink ratio, a trait highly conserved across environments. HSS were sink-limited under e-CO_2_, probably associated with low triose phosphate utilization (*TPU*) capacity. We suggest that the local C source-sink ratio is a potential target for selecting more CO_2_-responsive cultivars, pending validation for a broader genotypic spectrum and for field conditions.

**Highlight:** Rice local carbon source-sink ratio and sink plasticity can drive genotypic responses of leaf photosynthesis and plant production in a CO_2_ elevation context.

## Introduction

Over the last 200 years, atmospheric carbon dioxide concentration [CO_2_] dramatically increased, attaining 410 μmol mol^-1^ today. This represents an increase of 35% compared to the pre-industrial level (NOAA Mauna Loa Observatory, Hawaii). Global climate models predict that [CO_2_] will reach 600 to 700 μmol mol^-1^ by 2050 (IPCC 2016). Carbon dioxide is the main substrate for photosynthesis and it is expected to be one of the main climate change factors impacting plant development (Becklin *et al.*, 2017) and crop production (Rosenzweig *et al.*, 2014; Ko *et al.*, 2014; Kromdijk and Long, 2016). It is thus likely to contribute to the 70% increase in crop production aimed at by 2050.

As a major staple food, rice will play an important role in this context. Increasing its photosynthetic capacity is thus a challenge for increasing crop yield (Yoshida *et al.*, 2008; Yoshida and Horie, 2009; Makino, 2011; Simkin *et al.*, 2019) and the performance of cropping systems (Lawson *et al.*, 2012; Evans, 2013; Long *et al.*, 2015; Ort *et al.*, 2015; Foyer *et al.*, 2017; Nuccio *et al.*, 2017). Several studies reported inter and intra-specific variability in rice photosynthetic rate increase under elevated CO_2_ (e-CO_2_) (Makino and Mae, 1999; Adachi *et al.*, 2014; Zhu *et al.*, 2014; Wang *et al.*, 2015). However, an increase in photosynthesis under e-CO_2_ was not systematically associated with an increase in grain yield (Kobayashi *et al.*, 2006; Long *et al.*, 2006, 2015; Ainsworth, 2008; Leakey *et al.*, 2012; Wang *et al.*, 2015; Hasegawa *et al.*, 2016). One of the main hypotheses to explain this observation is that not only C source capacity but also C sink capacity drives crop production, and that either source or sink capacity alone can explain e-CO_2_ impacts on yield (Körner, 2015; Ludewig and Sonnewald, 2016; Burnett *et al.*, 2016; Sonnewald and Fernie, 2018).

It has been reported that a C source-sink imbalance characterized by a small sink (demand) combined with a large source capacity (supply) can down-regulate photosynthesis due to the accumulation of end-products in leaf photosynthetic tissues (Paul and Pellny, 2003; Ainsworth and Bush, 2011; Lemoine *et al.*, 2013; White *et al.*, 2016; Ruiz-Vera *et al.*, 2017). Accordingly, an increase in the photosynthetic rate, particularly if caused by e-CO_2_, should be accompanied with an increase in sink capacity. With this respect, phenotypic plasticity, the plant capacity to adjust its phenotype to environment including resources, may represent opportunities to improve crops for future environments (Franks *et al.*, 2014).

The C source-sink relationships affecting leaf photosynthetic rate are complex as they involve dynamic feedbacks among different levels of morphological, physiological and biochemical organization. Modelling is an integrative and quantitative way to analyze them (Poorter *et al.*, 2013; Wu *et al.*, 2016). Several models were developed to describe C source-sink interactions as reviewed by Chang and Zhu (2017), but models’ predictive power was found to be low, especially under e-CO_2_ (Asseng *et al.*, 2013). Most of these models are inspired by the mechanistic biochemical model of photosynthesis by Farquhar *et al.* (1980) (FvCB). FvCB is today used as a module within crop models. However, as it takes into account only partially the negative feedback of the accumulation of end-products on photosynthesis (Leegood and Furbank, 1986; Sharkey *et al.*, 1986), this model has limitation in dealing with C source-sink imbalanced situations. The feedback is partly controlled by the Triose Phosphate Utilization (*TPU*) parameter that sets a dynamic maximum leaf photosynthetic capacity (Sharkey *et al.*, 1986; Paul and Foyer, 2001). *TPU* has been less studied than other photosynthetic parameters. It acts at the interface between the production and the consumption of assimilates (Yang *et al.*, 2016) and may as such be considered when predicting photosynthesis and crop performance under e-CO_2_ conditions (Busch and Sage, 2016; Rogers Alistair *et al.*, 2016; Lombardozzi *et al.*, 2018; Busch *et al.*, 2018; McClain and Sharkey, 2019; Fabre *et al.*, 2019).

In a previous paper (Fabre *et al.*, 2019), we observed strong reductions in leaf photosynthesis in rice subjected to artificially induced sink limitation, through panicle pruning and/or elevated atmospheric CO_2_, particularly in the afternoon when sugar translocation probably became a bottleneck. While sucrose accumulated in the leaf, *TPU* and *V*_*cmax*_ decreased. There is limited information, however, on how genotypic variation in C source and sink capacity during grain filling (or their adaptive plasticity) affects the response of photosynthesis and yield to e-CO_2_. Genotypic variation in photosynthetic parameters and assimilate allocation patterns to sinks has been reported (Flood *et al.*, 2011; Gu *et al.*, 2012), but little is known on how these traits interact. Rice cultivars differ strongly in the biomass and grain yield response to e-CO_2_ in open-air FACE field trials (Hasegawa *et al.*, 2013; Wang *et al.*, 2015). The genotypic differences could in part be explained with panicle sink capacity (Hasegawa et al., 2013) or compensatory tillering ability (Kikuchi et al., 2017).

This paper aims to investigate genotypic variation in the response of photosynthesis and growth to e-CO_2_ conditions in rice cultivars, differing constitutively in morphological traits conferring source and sink capacity. We selected five *indica*-type rice cultivars differing in the local C source-sink ratio, characterized here using a proxy parameter, i.e., the ratio of flag leaf size to the grain number of the corresponding panicle. Plants were grown under e-CO_2_ (800 μmol mol^-1^) conditions either during grain filling or throughout the whole plant development cycle. The specific objectives were to (1) study the effect in controlled environments of e-CO_2_ (800 μmol mol^-1^) on photosynthetic parameters, yield and yield components; (2) observe levels of non-structural carbohydrates (NSC) in leaves and stem internodes; and (3) relate the findings to genotypic values of local C source-sink ratio. We will discuss the results with respect to opportunities to breed for traits that would enable a greater genotypic response to the ongoing and anticipated increase of atmospheric CO_2_ resources.

## Material and methods

### Plant material and growth conditions

We chose five high yielding *indica* rice cultivars (WAS182, TEQUING, WAS197, IR52, IR64) out of the 300 accessions in the PRAY *indica* panel (http://ricephenonetwork.irri.org/), based on their diversity in local C source-sink ratio (Table S1). As indicated above, the exact local C source-sink ratio is hard to quantify, we used the ratio of flag leaf area (source) to the fertile spikelet number of the corresponding panicle (sink) on the main stem, according to Fabre *et al.*, (2019). For this purpose, field phenomics data were provided by CIAT (Columbia) (Rebolledo, unpublished) for 2013 and 2014 experiments. In order to minimize others sources of variations, cultivars were selected with similar phenological and morphological traits such as degree days to flowering, tiller and panicle number per plant, plant height, and grain fertility (Table S1).

The five genotypes were germinated on wet filter paper and transplanted into 6L pots filled with EGO 140 substrate (17%N-10%P-14%K, pH = 5). This pot size is sufficient for rice to avoid reductions in photosynthesis or biomass accumulation along plant cycle (Sage, 1994; Poorter *et al.*, 2012). Basal fertilizer was applied before transplanting using Basacot 6 M (Compo Expert, France) at 2 g l^-1^, 11%N-9%P-19%K +2%Mg. A second application was performed just before heading stage to avoid post-floral nitrogen deficiency.

Plants were grown from April to August 2018 at the Agronomical Research and International Center for Development (CIRAD, Montpellier, France) in two adjacent, climate-controlled greenhouse compartments. They were grown under natural daylight with supplemental lighting maintaining a 12-h photoperiod using horticultural red-blue LED projectors (Alpheus Radiometrix 15M1006) providing an R/FR ratio of about 1.2. Microclimate was monitored using data-loggers (CR1000 Campbell Scientific) installed in each compartment. Air temperature (Ta) averaged 29 °C (day) and 22 °C (night) as measured with a PT1000 probe under fan-aspirated shield. Air relative humidity (RH) averaged 65% (day) and 80% (night), measured by HMP45 (Vaisala, Helsinki, Finland) and PPFD was measured with a SKP215 (Skye Instrument quantum sensor, Powys, UK) providing on average photosynthetic irradiance of 800 μmol m^-2^ s^-1^ at the top of the canopy level during the daytime. The mean photosynthetically active radiation received by the plants during their life cycle was 15.12 MJ m^-2^ d^-1^.

A total of 336 pots including border pots were arranged at 20-cm spacing among plants in a randomized design with five replications per cultivar and per CO_2_ treatment on movable tables. Pots were kept watered at field capacity while maintaining the perforated pot bottoms in 5 cm of standing water. To minimize border effects on each table, border plants on the tables were not used for measurements. The tables were moved weekly to avoid effects of spatial heterogeneity. We used two compartments that were differentiated only by the atmospheric CO_2_ level: 400 μmol mol^-1^ (ambient) in one compartment versus 800 μmol mol^-1^ (e-CO_2_) in the other.

Two complementary experiments (EXP1 and EXP2) were then carried out using independent sets of plants within the 336 plants available:

EXP1 was dedicated to a physiological characterization of photosynthesis response to variation in plant C source-sink balance during a medium-term (15-day) e-CO_2_ period from heading stage onwards. For this purpose, 25 plants (5 per cultivar) from ambient CO_2_ compartment were transferred to the e-CO_2_ compartment at heading stage, while 25 others remained in the ambient compartment. At this stage, the spikelet number per panicle is fixed on the main stem (Cock and Yoshida, 1973). The e-CO_2_ treatment was continued for a 15 days, corresponding to the grain filling period (Cho *et al.*, 1988). Applying e-CO_2_ during grain filling was chosen to ensure that the five genotypes differed only for their local C source-sink ratio, avoiding the appearance of new sinks (tillers) or the increase of existing ones (spikelets per panicle). Morphological, physiological and biochemical measurements were performed at the end of the 15-d treatment as detailed below.

In EXP2, the two CO_2_ treatments (ambient and e-CO_2_) were differentiated from transplanting to maturity, with 25 plants (5 per genotype) plus border plants placed in each compartment (CO_2_ treatment). For each treatment, plants were characterized for growth and development traits along the cycle, photosynthesis and biochemical measurements at 15 days after heading, final biomass and grain production as described hereafter.

### Leaf photosynthesis measurement

Photosynthesis measurements were performed *in situ* on the fully expanded flag leaf on the main stem of five plants per cultivar in each treatment, 2 weeks after heading in both experiments. Comparison between the CO_2_ treatments was made by using simultaneously two infrared gas analyzers (GFS-3100, Walz, Germany) identically calibrated. Leaf net photosynthetic rate at cuvette CO_2_ concentration at 400 μmol mol^-1^ (*A*_*400*_) was evaluated using saturating PPFD light of 1500 μmol m^-2^ s^-1^, controlled leaf temperature at 28°C, relative humidity in the cuvette set to 65%, and constant air flow rate through the cuvette of 800 ml min^-1^. A large exchange area cuvette of 8 cm^2^ was used to limit border effects known to affect photosynthesis measurement at high CO_2_ concentration (Long and Bernacchi, 2003). Simultaneously, chlorophyll fluorescence was measured using Walz PAM-fluorimeter 3055FL integrated to the photosynthesis equipment. The steady-state fluorescence yield (*F*_s_) was measured after registering the gas-exchange parameters. A saturating light pulse (8000 μmol m^-2^ s^-1^ during 0.8 s) was applied to achieve the light-adapted maximum fluorescence (*F*_m_*’*). The operating PSII photochemical efficiency (φPSII) was determined as φPSII = (*F*_m_’ – *F*_s_)/*F*_m_’. Because a photosynthesis down-regulation related to C sink limitation increases along the day (Fabre *et al.*, 2019), all measurements (EXP1 and EXP2) were made 6 hours after dawn.

Often, the maximum photosynthetic rate is determined by the rate of conversion and use of triose phosphates into starch and sucrose (Sharkey *et al.*, 1986; McClain and Sharkey, 2019). In EXP1 only, after the measurement for *A*_*400*_, the CO_2_ concentration in the measurement cuvette was increased through the following step-wise sequence: 900, 1300 and 1600 μmol mol^-1^, and waiting for steady-state photosynthesis at each step. This allowed to identify whether the *TPU* limitation occurred. We also used fluorescence data measured in combination to confirm any TPU limitation, as a decline in φPSII with increasing intercellular CO_2_ concentration (*C*_i_) would suggest a TPU limitation (Sharkey, 2016). The leaf photosynthesis rate under saturating light and CO_2_ levels (1600 μmol mol^-1^) was taken as the maximum leaf photosynthesis capacity level (*Amax*) and this was measured in both CO_2_ treatments. Then, the ratio of the average *Amax* of plants grown at e-CO_2_ by the average *Amax* of plants grown at ambient CO_2_ (*Amax*_800_/*Amax*_400_) was calculated.

### Sugar content analysis

Immediately after photosynthesis measurement, the same leaf was sampled to determine non-structural carbohydrate content (NSC: starch, sucrose, glucose, and fructose). Prior to grinding with a ball grinder (Mixer mill MM 200, Retsch, Germany), the samples were frozen in liquid nitrogen. The sugars were extracted three times from 20 mg samples with 1 mL of 80% ethanol for 30 min at 75°C, and then centrifuged for 10 min at 9500 g (Mikro 200, Hettich centrifuge). Soluble sugars (sucrose, glucose, and fructose) were contained in the supernatant and starch in the sediment. The supernatant was filtered in the presence of polyvinyl polypyrrolidone and activated carbon to eliminate pigments and polyphenols. After evaporation of solute with Speedvac (RC 1022 and RCT 90, Jouan SA, Saint Herblain, France), soluble sugars were quantified by high performance ionic chromatography (HPIC, standard Dionex) with pulsed amperometric detection (HPAE-PAD). The sediment was solubilized with 0.02 N NaOH at 90° C for 1 h 30 min and then hydrolyzed with a-amyloglucosidase at 50 °C, pH 4.2 for 1 h 30 min. Starch was quantified as described by Boehringer (Pomeranz and Meloan, 1994) with 5 μL of hexokinase (glucose-6-phosphate dehydrogenase), followed by spectro-photometry of NADPH at 340 nm (spectrophotometer UV/VIS V-530, Jasco Corporation, Tokyo, Japan).

### Leaf Nitrogen and Potassium Content

In EXP1 only, the leaf blade situated below the flag leaf used for gas exchange measurements was analyzed for nitrogen and potassium concentration (expressed in percent of the leaf blade dry matter: Nm or Km [mg g^-1^]), specific leaf area (SLA [cm^2^ g^-1^]), and on this basis, nitrogen and potassium content on leaf-area basis by multiplying Nm or Km by SLA^-1^ (Na or Ka [g N m^-2^]). The area of each leaf was measured with a leaf area meter (LI-3100 LI-COR, Lincoln, NE, USA) and the leaf then oven-dried until constant weight (48 h at 70 °C). Total leaf N was analyzed based on Dumas combustion method using a LECO TruMac Nitrogen analyzer, and K content was measured using an ICP-OES 700 series spectrometer (Agilent Technologies). A relative indicator of chlorophyll content (SPAD) was measured on the same leaf using a SPAD-502 (Minolta, Ltd., Japan) in both experiments.

### Plant growth, biomass and yield component measurements

After samplings for biochemical analyses performed in EXP1, plant shoots were harvested. Tillers and panicles were counted and total plant green leaf area measured, using a leaf area meter (LI-3100 LI-COR). Then, total stem and leaf dry matter per plant (DM) were measured separately after drying at 70°C during 48h and adding DM from organs used for biochemical analyses. Plant height was determined as the vertical distance from the root-shoot junction to the ligule of the flag leaf.

In EXP2, tiller and panicle number, plant height, plant green leaf area and DM were measured at maturity stage as described. Total plant shoot DM (stems + leaves) and plant bulk SLA (cm^2^ g^-1^) were calculated on this basis. Grain yield and yield components were calculated according to (Liu *et al.*, 2008): grains were sorted by using a densiometric column and the number of filled (fertile) and empty grains was counted using a grain counter to determine the filled spikelet percentage. Dry weight of ripened grains was determined after oven-drying at 80°C for 72 h. The 1000-grains dry weight was then calculated. Grain yield per m^2^ was computed by multiplying grain production per plant in g by the number of plants per m^2^.

In both experiments, local C source-sink ratio was estimated for each plant by dividing flag leaf blade area and fertile spikelet number on the main stem.

### Statistical analysis

Physiological, biochemical traits and yield components were analyzed as a completely randomized design using a two-way analysis of variance of CO_2_ treatment, genotype and interaction using R (version 3.5.2, R Foundation for Statistical Computing), after testing for normal distribution. Wherever appropriate, comparison between means was performed using Tukey *post-hoc* test (α=0.05) with the same software. In the absence of normal distribution (e.g. carbohydrate), data were log-transformed to stabilize the variance. Principal component analysis (PCA) was performed with the same software using FactoMineR package to analyze covariation.

## Results

### Local C source-sink ratio

In both experiments, a highly significant cultivar effect (P<0.001) was noticed for the local C source-sink ratio on the main stem (Tables 1 & 2) with neither CO_2_ treatment effects nor interaction effects. The genotypic differences were conserved between EXP1 and EXP2 as indicated by the linear correlation between this ratio in medium vs. long term e-CO_2_ treatments (Fig. S1; R^2^=0.99, P<0.01).

**Table 1:** Summary of a 2-way analysis of variance for growth, biochemical and physiological traits measured two weeks after heading on five rice cultivars grown under two [CO_2_] levels (EXP1 : 400 μmol mol^-1^ continuously and 800 μmol mol^-1^ during 15 days from heading). Average values ± SE (n=5) are presented. For each column, values followed by different letters differ significantly (P<0.05) *** P<0.001, ** P<0.01, * P<0.05, ns – not significant

| Cultivar | [CO<sub>2</sub>]<br>grown | Tiller number<br>per plant | Panicle<br>number per<br>plant | Plant height<br>(cm) | Stem DM<br>(g plant<sup>-1</sup>) | Leaf DM<br>(g plant<sup>-1</sup>) | Flag leaf N<br>content<br>(g m<sup>-2</sup>) | Flag leaf K<br>content<br>(g m<sup>-2</sup>) | Main stem<br>panicle spikelets<br>number (SP) | Main stem<br>flag leaf area<br>(FLA) (cm<sup>2</sup>) | Main stem C<br>source-sink<br>ratio (FLA/SP) | Flag leaf<br>specific area<br>SLA (cm<sup>2</sup> g<sup>-1</sup>) | SPAD |
| --- | --- | --- | --- | --- | --- | --- | --- | --- | --- | --- | --- | --- | --- |
| WAS182 | 400 | 13.0 ± 0.3 a | 10.5 ± 0.2 a | 85.8 ± 1.7 a | 36.8 ± 0.6 ab | 14.1 ± 0.7 a | 1.5 ± 0.1 a | 1.1 ± 0.1 a | 231.4 ± 11.0 a | 40.7 ± 3.9 bc | 0.17 ± 0.01 de | 151.3 ± 9.3 a | 42.4 ± 0.3 a |
|  | 800 | 12.8 ± 0.3 a | 10.8 ± 0.3 a | 85.4 ± 1.1 a | 39.3 ± 1.2 a | 14.3 ± 0.7 a | 1.5 ± 0.2 a | 1.1 ± 0.0 a | 238.2 ± 8.9 a | 41.4 ± 2.3 bc | 0.17 ± 0.01de | 144.9 ± 17.7 a | 43.3 ± 0.6 a |
| TEQUING | 400 | 13.0 ± 0.7 a | 10.4 ± 0.7 a | 85.6 ± 2.2 a | 38.4 ± 2.2 ab | 12.3 ± 0.8 ab | 1.4 ± 0.0 a | 0.9 ± 0.0 a | 247.4 ± 10.1 a | 39.0 ± 1.2 c | 0.15 ± 0.00 e | 159.8 ± 6.0 a | 44.3 ± 0.8 a |
|  | 800 | 13.0 ± 0.4 a | 10.4 ± 0.5 a | 84.6 ± 0.8 a | 39.3 ± 0.4 a | 11.9 ± 0.5 ab | 1.3 ± 0.1 a | 1.0 ± 0.0 a | 236.2 ± 5.8 a | 40.1 ± 4.0 c | 0.17 ± 0.01 de | 157.5 ± 19.6 a | 43.5 ± 0.6 a |
| WAS197 | 400 | 13.6 ± 0.5 a | 11.4 ± 0.4 a | 83.2 ± 0.5 a | 29.1 ± 1.9 bc | 11.7 ± 0.6 ab | 1.3 ± 0.0 a | 0.9 ± 0.0 a | 215.4 ± 8.6 ab | 47.5 ± 2.4 abc | 0.22 ± 0.00 cde | 180.2 ± 12.8 a | 43.0 ± 0.6 a |
|  | 800 | 13.8 ± 0.4 a | 11.2 ± 0.2 a | 84.8 ± 0.9 a | 31.5 ± 0.9 abc | 11.5 ± 0.7 ab | 1.2 ± 0.0 a | 1.0 ± 0.1 a | 224.0 ± 4.4 ab | 50.8 ± 3.6 abc | 0.22 ± 0.01 cd | 182.6 ± 9.2 a | 42.9 ± 0.5 a |
| IR52 | 400 | 12.8 ± 0.3 a | 10.6 ± 0.2 a | 83.0 ± 1.3 a | 33.6 ± 2.0 abc | 12.6 ± 0.7 ab | 1.5 ± 0.1 a | 1.1 ± 0.0 a | 190.4 ± 7.5 bc | 55.4 ± 4.3 ab | 0.29 ± 0.01 ab | 147.3 ± 4.6 a | 44.2 ± 0.3 a |
|  | 800 | 14.0 ± 0.3 a | 10.6 ± 0.6 a | 82.0 ± 0.8 a | 32.7 ± 3.8 abc | 12.6 ± 0.7 ab | 1.3 ± 0.1 a | 1.1 ± 0.1 a | 188.4 ± 10.4 bc | 56.7 ± 1.2 a | 0.30 ± 0.01 a | 156.9 ± 15.9 a | 43.5 ± 0.7 a |
| IR64 | 400 | 14.0 ± 0.7 a | 11.6 ± 0.5 a | 80.0 ± 2.1 a | 26.5 ± 2.4 c | 10.2 ± 0.8 b | 1.6 ± 0.1 a | 1.0 ± 0.0 a | 167.0 ± 12.0 c | 38.8 ± 3.7 c | 0.23 ± 0.01 bcd | 176.3 ± 6.1 a | 42.6 ± 0.4 a |
|  | 800 | 13.8 ± 0.7 a | 11.6 ± 0.5 a | 79.8 ± 0.7 a | 26.4 ± 1.9 c | 10.0 ± 0.5 b | 1.5 ± 0.1 a | 1.0 ± 0.1 a | 152.8 ± 7.5 c | 39.4 ± 2.0 c | 0.25 ± 0.01 abc | 188.4 ± 11.5 a | 42.8 ± 0.4 a |
| [CO<sub>2</sub>] |  | ns | ns | ns | ns | ns | ns | ns | ns | ns | ns | ns | ns |
| Cultivar |  | ns | ns | ** | *** | *** | ns | ns | *** | *** | *** | * | ns |
| [CO<sub>2</sub>]x Cultivar |  | ns | ns | ns | ns | ns | ns | ns | ns | ns | ns | ns | ns |
| Cultivar | [CO<sub>2</sub>]<br>grown | A<sub>400</sub><br>(μmol m<sup>-2</sup> s<sup>-1</sup>) | C<sub>i</sub> ambient<br>(μmol mol<sup>-1</sup>) | A<sub>max</sub><br>(μmol m<sup>-2</sup> s<sup>-1</sup>) | C<sub>i</sub> elevated<br>(μmol mol<sup>-1</sup>) | Ratio<br>A<sub>max800</sub>/A<sub>max400</sub> | Sucrose flag leaf<br>(μg cm<sup>-2</sup>) | Sucrose<br>top internode<br>(mg g<sup>-1</sup> DM) | Hexose flag leaf<br>(μg cm<sup>-2</sup>) | Hexose<br>top internode<br>(mg g<sup>-1</sup> DM) | Starch flag leaf<br>(μg cm<sup>-2</sup>) | Starch<br>top internode<br>(mg g<sup>-1</sup> DM) |  |
| WAS182 | 400 | 19.6 ± 0.3 d | 283.7 ± 2.0 a | 24.2 ± 0.6 bc | 1021.0 ± 41.6 a | 1.20 ± 0.08 bc | 310.3 ± 15.6 ab | 209.2 ± 11.4 ab | 10.65 ± 0.3 ab | 49.3 ± 8.6 a | 40.1 ± 8 abc | 136.3 ± 17.3 c |  |
|  | 800 | 24.5 ± 0.5 a | 267.4 ± 5.6 a | 29.2 ± 1.6 a | 1004.4 ± 12.4 a |  | 271.6 ± 13.1 ab | 227.3 ± 19 a | 11.94 ± 0.7 a | 41.3 ± 11.8 a | 51.3 ± 9.7 ab | 187.6 ± 24.9 bc |  |
| TEQUING | 400 | 17.3 ± 0.4 e | 287.4 ± 6.4 a | 21.6 ± 0.5 c | 1082.4 ± 39.6 a | 1.34 ± 0.07 c | 336.2 ± 6.4 a | 127 ± 5.7 de | 13.03 ± 0.5 a | 8.9 ± 1.2 bc | 35.8 ± 4.4 abc | 340.3 ± 14.6 ab |  |
|  | 800 | 21.0 ± 0.1 cd | 288.4 ± 2.6 a | 29.1 ± 1.2 a | 1059.0 ± 17.6 a |  | 347.6 ± 28.8 a | 139.1 ± 20.1 e | 13.81 ± 1.2 a | 9.3 ± 4.8 c | 61.2 ± 5.2 a | 383.1 ± 24.5 a |  |
| WAS197 | 400 | 21.5 ± 0.4 bc | 289.1 ± 9.1 a | 25.0 ± 0.6 abc | 1061.2 ± 21.6 a | 1.10 ± 0.03 ab | 276.2 ± 20.9 ab | 194 ± 7.9 abc | 15.92 ± 2.3 a | 63 ± 5.6 a | 43.1 ± 6.6 ab | 143.3 ± 21.6 c |  |
|  | 800 | 23.2 ± 0.3 ab | 289.9 ± 6.2 a | 27.4 ± 0.7 ab | 1117.3 ± 20.0 a |  | 246.1 ± 17.2 abc | 198 ± 6.3 abc | 11.82 ± 1.4 a | 43 ± 5.1 a | 47.6 ± 3.9 ab | 185.9 ± 12 abc |  |
| IR52 | 400 | 19.8 ± 0.2 cd | 264.2 ± 6.0 a | 23.5 ± 0.8 bc | 1144.4 ± 19.1 a | 0.90 ± 0.03 a | 191.8 ± 13.6 c | 157.7 ± 8.6 bcde | 6.73 ± 1 b | 26.4 ± 3.7 ab | 18.6 ± 1.2 c | 157.8 ± 13.1 c |  |
|  | 800 | 17.0 ± 0.4 e | 265.5 ± 3.4 a | 21.2 ± 0.9 c | 1148.1 ± 43.9 a |  | 255.3 ± 23.2 abc | 153.2 ± 4.3 cde | 9.75 ± 1.6 ab | 38.4 ± 3.2 a | 30.4 ± 5.3 bc | 145.2 ± 37.5 c |  |
| IR64 | 400 | 19.3 ± 0.4 d | 271.6 ± 5.6 a | 25.3 ± 0.7 abc | 1046.5 ± 32.6 a | 0.97 ± 0.03 ab | 221.4 ± 17.8 bc | 194.1 ± 7.1 abc | 9.94 ± 1.1 ab | 23.4 ± 3.1 ab | 20.5 ± 5.1 c | 156.2 ± 18.9 c |  |
|  | 800 | 19.7 ± 0.5 cd | 266.9 ± 4.6 a | 24.8 ± 0.9 bc | 1052.6 ± 55.0 a |  | 274.7 ± 20.8 ab | 182.7 ± 7.4 abcd | 12.17 ± 1.7 a | 23.2 ± 4.3 ab | 35.2 ± 7.2 bc | 154.6 ± 24.1 c |  |
| [CO<sub>2</sub>] |  | *** | ns | *** | ns | / | ns | ns | ns | ns | *** | ns |  |
| Cultivar |  | *** | ** | *** | ** | *** | *** | *** | *** | *** | *** | *** |  |
| [CO<sub>2</sub>]x Cultivar |  | *** | ns | *** | ns | / | * | ns | ns | ns | ns | ns |  |

**Table 2:** Summary of a 2-way analysis of variance for biochemical and physiological parameters measured two weeks after heading on the flag leaf on the main stem, and plant growth and yield components evaluated at maturity on five rice cultivars grown under two [CO_2_] levels (EXP2: 400 vs 800 μmol mol^-1^ continuously from seedling to maturity). Average values ± SE (n=5) are presented. For each column, values followed by different letters differ significantly (P<0.05) *** P<0.001, ** P<0.01, * P<0.05, ns – not significant

| Cultivar | [CO<sub>2</sub>]<br>grown | Tiller number per<br>plant | Panicle number<br>per plant | Plant height<br>(cm) | total shoot DM<br>(g) | 1000-grains weight<br>(g) | Grain Yield<br>(g m<sup>-2</sup>) | Filled spikelet<br>ratio (%) | Main stem panicle<br>spikelets number<br>(SP) | Main stem<br>flag leaf area<br>(FLA) (cm<sup>2</sup>) | Main stem C<br>source-sink ratio<br>(FLA/SP) |
| --- | --- | --- | --- | --- | --- | --- | --- | --- | --- | --- | --- |
| WAS182 | 400 | 12.0 ± 0.3 a | 10.4 ± 0.6 a | 81.0 ± 0.6 a | 50.3 ± 1.1 abc | 21.9 ± 0.0 a | 598.8 ± 18.1 ab | 70.9 ± 3.5 b | 234.4 ± 7.4 abc | 36.2 ± 4.0 c | 0.15 ± 0.02 c |
| | 800 | 14.0 ± 0.3 a | 12.2 ± 0.4 a | 85.7 ± 1.6 a | 59.1 ± 2.5 ab | 22.7 ± 0.0 a | 666.2 ± 13.9 a | 75.4 ± 2.1 ab | 247.2 ± 8.8 ab | 41.8 ± 1.7 abc | 0.17 ± 0.00 c |
| TEQUING | 400 | 12.4 ± 0.2 a | 11.4 ± 0.4 a | 84.7 ± 1.1 a | 41.5 ± 1.6 c | 22.1 ± 0.3 a | 570.9 ± 20.7 abc | 86.6 ± 1.8 a | 235.6 ± 8.4 abc | 38.7 ± 4.7 bc | 0.16 ± 0.02 c |
| | 800 | 13.8 ± 0.6 a | 12.8 ± 0.8 a | 83.8 ± 1.6 a | 54.0 ± 3.0 ab | 22.8 ± 0.4 a | 657.8 ± 13.9 a | 90.1 ± 1.4 a | 261.8 ± 6.0 a | 46.1 ± 3.2 abc | 0.18 ± 0.01 c |
| WAS197 | 400 | 13.2 ± 0.9 a | 12.8 ± 0.8 a | 82.0 ± 1.9 a | 39.9 ± 1.7 c | 22.3 ± 0.1 a | 610.2 ± 39.6 a | 66.7 ± 1.0 b | 203.4 ± 6.7 cde | 41.6 ± 1.3 abc | 0.21 ± 0.01 bc |
| | 800 | 14.0 ± 0.5 a | 13.0 ± 0.4 a | 82.2 ± 1.1 a | 48.2 ± 1.3 abc | 22.8 ± 0.2 a | 656.7 ± 29.6 a | 68.0 ± 2.0 b | 218.6 ± 4.7 bcd | 45.8 ± 2.0 abc | 0.21 ± 0.01 bc |
| IR52 | 400 | 11.4 ± 0.5 a | 11.4 ± 0.5 a | 83.0 ± 1.5 a | 57.4 ± 2.3 ab | 22.5 ± 0.2 a | 548.5 ± 7.9 abc | 68.5 ± 1.3 b | 184.4 ± 9.6 ef | 53.1 ± 3.1 ab | 0.29 ± 0.02 a |
| | 800 | 12.4 ± 0.9 a | 11.8 ± 1.1 a | 85.2 ± 0.9 a | 60.4 ± 2.7 a | 22.9 ± 0.0 a | 573.1 ± 26.1 abc | 70.6 ± 4.8 b | 191.2 ± 6.2 def | 53.2 ± 3.3 a | 0.28 ± 0.01 a |
| IR64 | 400 | 13.2 ± 0.5 a | 12.6 ± 0.5 a | 79.4 ± 0.9 a | 46.3 ± 2.9 bc | 23.2 ± 0.3 a | 412.2 ± 23.5 c | 65.9 ± 1.3 b | 159.0 ± 4.6 f | 39.0 ± 3.0 abc | 0.24 ± 0.01 ab |
| | 800 | 14.0 ± 0.3 a | 13.0 ± 1.0 a | 80.0 ± 0.9 a | 50.6 ± 3.4 abc | 23.1 ± 0.4 a | 426.3 ± 27.1 bc | 65.6 ± 2.9 b | 164.6 ± 4.3 f | 38.3 ± 1.2 bc | 0.23 ± 0.01 ab |
| [CO<sub>2</sub>] |  | ** | ns | ns | *** | ns | ** | ns | ** | ns | ns |
| Cultivar |  | * | ns | * | *** | ns | *** | *** | *** | *** | *** |
| [CO<sub>2</sub>]×Cultivar | | ns | ns | ns | ns | ns | ns | ns | ns | ns | ns |
| Cultivar | [CO<sub>2</sub>]<br>grown | A<sub>400</sub><br>($\mu\text{mol m}^{-2} \text{s}^{-1}$) | C<sub>i</sub> ambient<br>($\mu\text{mol mol}^{-1}$) | Sucrose flag leaf<br>($\mu\text{g cm}^{-2}$) | Sucrose<br>top internode<br>(mg g<sup>-1</sup> DM) | Hexose flag leaf<br>($\mu\text{g cm}^{-2}$) | Hexose<br>top internode<br>(mg g<sup>-1</sup> DM) | Starch flag leaf<br>($\mu\text{g cm}^{-2}$) | Starch<br>top internode<br>(mg g<sup>-1</sup> DM) | SPAD | Plant SLA<br>(cm<sup>2</sup> g<sup>-1</sup>) |
| WAS182 | 400 | 19.6 ± 0.3 bcd | 283.7 ± 2.0 a | 310.3 ± 15.6 a | 209.2 ± 11.4 ab | 10.65 ± 0.3 abc | 49.3 ± 8.6 ab | 40.1 ± 8 abc | 136.3 ± 17.3 c | 42.2 ± 0.4 a | 220.2 ± 7.1 a |
| | 800 | 22.6 ± 0.3 a | 277.9 ± 11.2 a | 297.4 ± 19.2 ab | 245.4 ± 18.3 a | 14.1 ± 0.8 ab | 46 ± 6.5 ab | 36.3 ± 4.3 bc | 182.2 ± 9.8 bc | 41.6 ± 0.6 a | 222.4 ± 14.3 a |
| TEQUING | 400 | 17.3 ± 0.4 d | 287.4 ± 6.4 a | 336.2 ± 6.4 a | 127 ± 5.7 c | 13.03 ± 0.5 ab | 8.9 ± 1.2 d | 35.8 ± 4.4 abc | 340.3 ± 14.6 ab | 44.3 ± 0.8 a | 221.5 ± 5.6 a |
| | 800 | 20.9 ± 0.4 abc | 274.3 ± 8.3 a | 344.1 ± 9.4 a | 116.3 ± 7.3 c | 18.5 ± 2.1 a | 8.3 ± 1.3 d | 61.9 ± 10.9 ab | 381 ± 19.3 a | 43.0 ± 0.4 a | 234.5 ± 8.7 a |
| WAS197 | 400 | 21.5 ± 0.4 ab | 297.0 ± 5.4 a | 276.2 ± 20.9 ab | 194 ± 7.9 ab | 15.92 ± 2.3 ab | 63 ± 5.6 a | 43.1 ± 6.6 abc | 143.3 ± 21.6 c | 43.0 ± 0.6 a | 259.3 ± 6.2 a |
| | 800 | 22.4 ± 1.1 a | 283.1 ± 10.2 a | 289.3 ± 24.6 ab | 182.2 ± 9.7 ab | 15.1 ± 1.6 ab | 55.1 ± 8.5 a | 75.2 ± 14.4 a | 152.3 ± 19.2 c | 42.9 ± 0.5 a | 217.9 ± 9.1 a |
| IR52 | 400 | 19.8 ± 0.2 abcd | 264.2 ± 6.0 a | 191.8 ± 13.6 c | 157.7 ± 8.6 bc | 6.73 ± 1 c | 26.4 ± 3.7 bc | 18.6 ± 1.2 c | 157.8 ± 13.1 c | 44.2 ± 0.3 a | 214.8 ± 5.2 a |
| | 800 | 18.5 ± 0.3 cd | 262.1 ± 6.0 a | 267.2 ± 22.5 abc | 155 ± 15.7 bc | 11.7 ± 0.9 ab | 30.1 ± 5.2 abc | 39.6 ± 11 abc | 108.3 ± 10.7 c | 43.1 ± 0.3 a | 221.3 ± 13.0 a |
| IR64 | 400 | 19.3 ± 0.4 bcd | 267.9 ± 4.9 a | 221.4 ± 17.8 bc | 194.1 ± 7.1 ab | 9.94 ± 1.1 bc | 23.4 ± 3.1 c | 20.5 ± 5.1 c | 156.2 ± 18.9 c | 42.6 ± 0.4 a | 247.3 ± 3.7 a |
| | 800 | 20.4 ± 0.6 abc | 263.3 ± 4.3 a | 258.2 ± 19.9 abc | 191.8 ± 12.5 ab | 12.4 ± 1.9 ab | 23.1 ± 3.4 c | 24.2 ± 5.1 c | 160.1 ± 30.3 c | 42.0 ± 0.7 a | 217.9 ± 7.9 a |
| [CO<sub>2</sub>] |  | *** | ns | * | ns | *** | ns | ** | ns | ns | ns |
| Cultivar |  | *** | ** | *** | *** | *** | *** | *** | *** | * | ns |
| [CO<sub>2</sub>]×Cultivar | | ** | ns | ns | ns | ns | ns | ns | ns | ns | * |

The five genotypes varied in local source-sink ratio, whereby two groups having the most contrasting values were significantly different in each experiment (Tukey’s test, Tables 1 & 2): the low source-sink ratio cultivars (LSS) WAS182 and TEQUING (small flag leaf area, large panicle sink potential) and the high source-sink ratio cultivar (HSS) IR52 (large flag leaf area, smaller panicle sink potential). WAS197 and IR64 showed intermediate values. A cultivar effect was also observed for flag leaf area (P<0.001) and spikelet number per panicle (P<0.001) on the main stem in both experiments, associated with a CO_2_ treatment effect in EXP2 for spikelet number (P<0.01). The genotypic variability in the local source-sink ratio was explained more by variation in spikelet number (36%) than by flag leaf area (9%). In EXP2, the CO_2_ treatment effect enhanced spikelet number in LSS cultivars by +8.3% on average (versus +3.8% in HSS). It enhanced flag leaf area by +17.2% for LSS (vs. +0.2% in HSS). This result suggests a greater plasticity of these traits for LSS cultivars in response to e-CO_2_.

### Photosynthetic parameters

Strong cultivar, CO_2_ and interaction effects (P<0.001, except interaction in EXP2 at P<0.01) were observed in both experiments for leaf net assimilation rate (*A*_*400*_) (Tables 1 and 2). This was mainly due to the significant increase in *A*_*400*_ for WAS182 and TEQUING cultivars (+23% in average in EXP1 and +17% in EXP2), whereas *A*_*400*_ was reduced in e-CO_2_ treatments for the HSS cultivar IR52 by 14% in EXP1 and 7% in EXP2 (Fig. 1A, B).

**Fig. 1:**
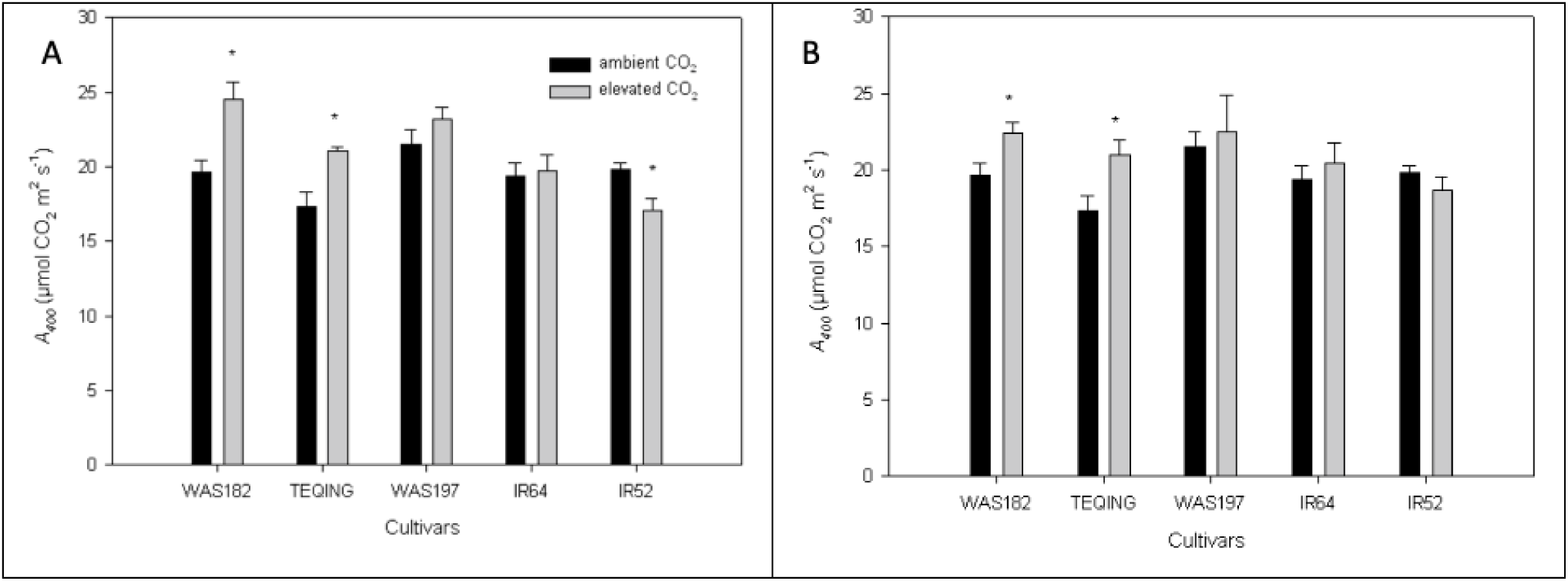
Net photosynthetic rate (*A*_*400*_) of the flag leaf on the main stem of five rice cultivars grown at two CO_2_ levels: ambient (400 μmol mol^-1^) and elevated (800 μmol mol^-1^) in two regulated greenhouse compartments. Measurements were taken 15 days after heading. Stars indicate significant differences at *p* < 0.05 (Tukey HSD test). Each bar represents the mean of 5 replicates ± SE. Panel **(A)** correspond to **EXP1** after 15 days CO_2_ enrichment from heading (medium-term) Panel **(B)** correspond to **EXP2** for an entire cycle CO_2_ enrichment (long-term)

Because stomatal conductance can affect *A* by changing CO_2_ intercellular concentration (*C*_i_), the photosynthesis of plants grown in ambient and e-CO_2_ should be compared at a same *C*_i_ to detect acclimation effects and to distinguish stomatal from non-stomatal effects. Here, variations in *A* were not associated with any significant variation of CO_2_ intercellular concentration despite a cultivar effect (P<0.01) (Table 1,2). This suggests that the photosynthetic capacity of the leaf *per se* was affected. No significant variations were observed on leaf chlorophyll content (SPAD) despite a small cultivars effect (P<0.05) only in EXP2.

Maximum photosynthetic capacity (*Amax*) was measured only in EXP1 (Table 1). Strong cultivar, CO_2_ and interaction effects were observed (P<0.001). A significant, positive response of *Amax* to CO_2_ enrichment was observed in LSS cultivars only (+20% for WAS182 and +34% for TEQUING) whereas a decrease was observed for the HSS cultivar IR52 (−10%) and IR 64 (−3%). These variations in photosynthetic capacity were not associated with differences in intercellular CO_2_ concentration (Table 1).

A negative and significant linear correlation (R^2^=0.90, P<0.05) was observed between *Amax* response to e-CO_2_ treatment (*Amax* ratio, *Amax*_800_/*Amax*_400_) and local source-sink ratio (Fig. 2). Under the assumption that *Amax* mainly reflects leaf *TPU* capacity (McClain and Sharkey, 2019), these results suggest that *TPU* capacity varied with genotypic, local C source-sink ratio. This was supported by a significant cultivar effect on *Amax* ratio (P<0.001, Table 1), indicating a significant enhancement in photosynthetic capacity for LSS cultivars under e-CO_2_.

**Fig. 2:**
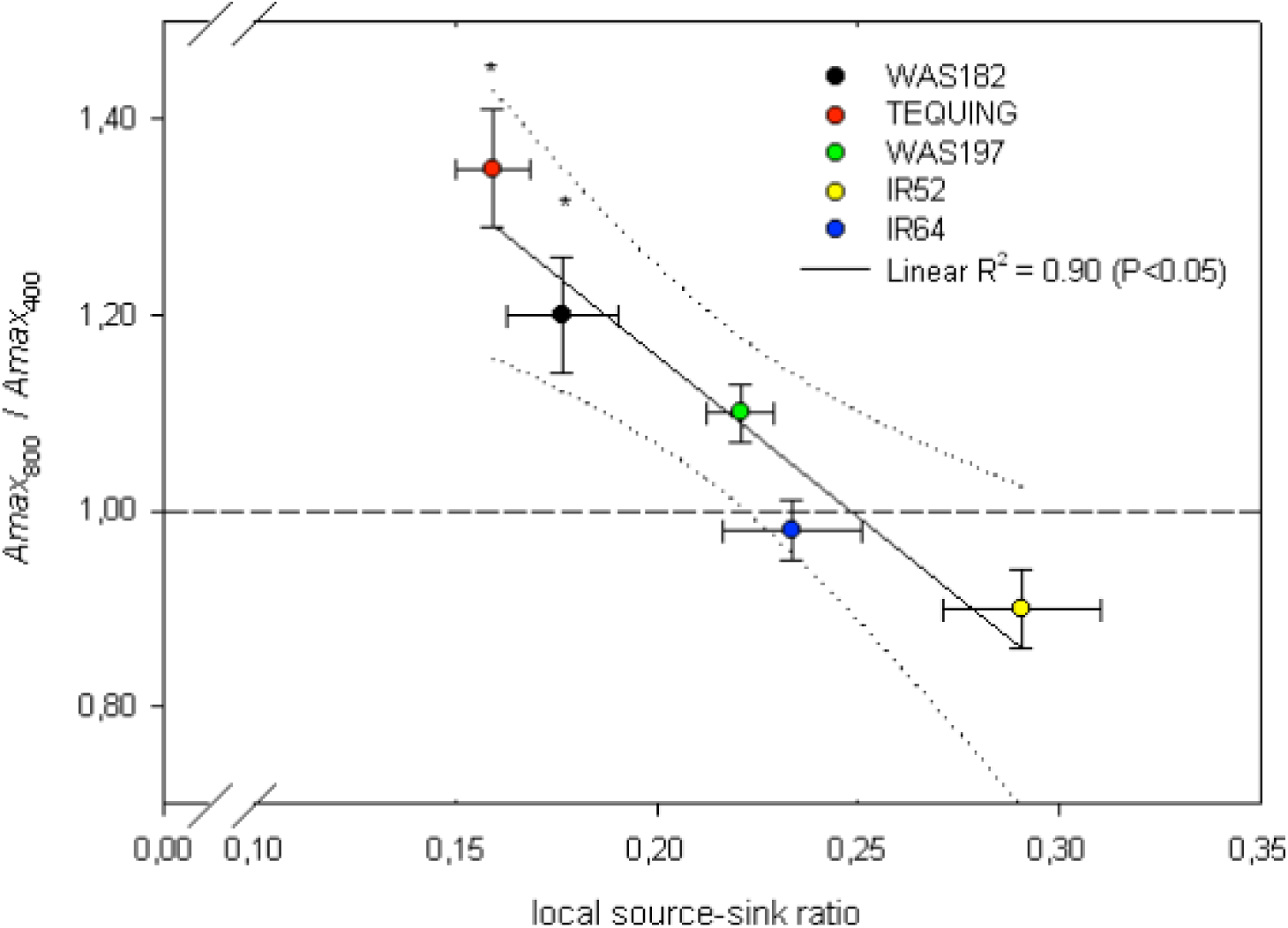
Relationship between *Amax*_800_/*Amax*_400_ (ratio between photosynthetic capacity (*Amax*) at elevated (800 μmol mol^-1^ [CO_2_]) and ambient (average value over 5 replicates at 400 μmol mol^-1^), versus local C source-sink ratio measured at ambient [CO_2_]. Datas from **EXP1** (15-day CO_2_ enrichment from heading) for the five studied rice cultivars were used. Each point is an average per genotype of 5 replicates ± SE. Stars indicate significant difference at *p* < 0.05 (Tukey HSD test) compared to ambient (dashed line).

Fluorescence measurements were performed concomitantly to photosynthesis measurements in order to assess the efficiency of PSII electron flow (φPSII) at three cuvette [CO_2_] levels and determine at which *C*_i_ level *TPU* became limiting for photosynthesis. In the case of HSS cultivar IR52 and intermediate cultivar IR64, a decline of φPSII was observed above a *C*_i_ of 680 μmol mol^-1^ on average highlighting a *TPU* limitation (Fig. 3B) under e-CO_2_ condition. However, as *C*_i_ thresholds for TPU limitation in terms of φPSII were higher than the *C*_i_ observed at *A*_*400*_ measurement; *TPU* did not limit *A*_*400*_ in any of the cultivars. A decline of φPSII at high cuvette [CO_2_] was not observed in LSS cultivars, whereby φPSII values were stable under e-CO_2_ condition (Fig. 3B).

**Fig. 3:**
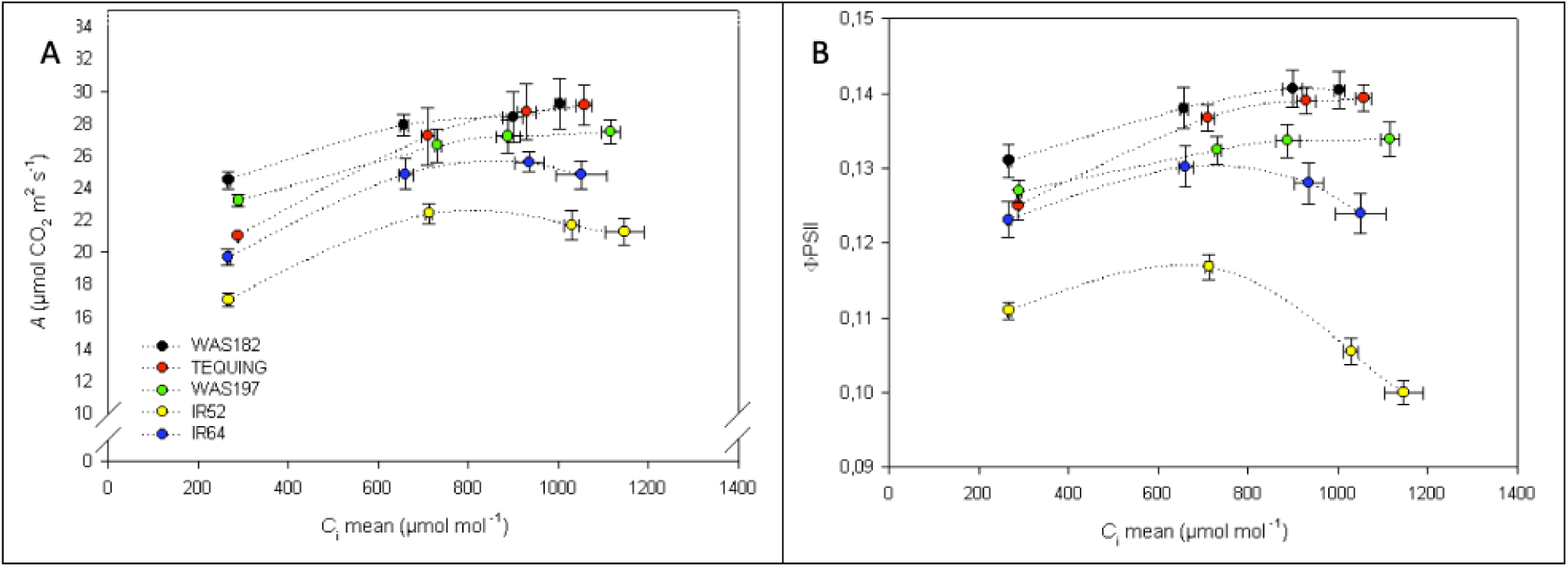
*A*/*C*_i_ (panel A) and corresponding φPSII/*C*_i_ curves (panel B) of five rice cultivars for EXP1, grown at 800 μmol mol^-1^. Measurements were taken 15 days after heading. Each point represents the mean of 5 replicates ± SE

### Plant growth and yield components

Overall, cultivars presented similar thermal time to flowering ranging from 1271 (WAS182) to 1373-degree days (IR64), with CO_2_ effects being absent (data not shown).

In EXP1, no CO_2_ treatment effect was observed on tiller and panicle numbers 15 days after heading. A cultivar effect was observed on plant height, stem DM and leaf DM (P<0.001, Table 1). LSS cultivars exhibited greater stem and leaf DM although differences were not significant. Flag leaf SLA only exhibited a slight cultivar effect (P<0.05, Table 1). No cultivar or treatment effects were observed for N and K content measured on the leaf below the flag leaf.

In EXP2, significant cultivar and CO_2_ effects, but no interactions, were observed for final shoot DM (P<0.001), tiller number (P<0.01) and grain yield (P<0.01, Table 2). e-CO_2_ enhanced plant shoot DM at maturity by 23% on average in LSS (significantly for TEQUING only), and only by 5% in the HSS cultivar. This could be partly explained by a stronger increase under e-CO_2_ of fertile tiller number at maturity in LSS cultivars (+15% on average) than in the HSS cultivar (3%). During the vegetative stage (45 days after emergence), maximum tiller number was increased by +34% on average for LSS cultivars, vs. +11% for HSS (Fig. S2).

Grain yield increase by e-CO_2_ differed among cultivars (13% on average in LSS, only 4% in HSS, Table 2). This could not be related to any variation in 1000-grain DM or filled grain percentage, although these yield components showed a slightly higher increase under e-CO_2_ in LSS cultivars (+3.5% averaged) compared to the HSS cultivar (1.5%). A significant cultivar effect was however observed on filled spikelet ratio (P<0.001, Table 2), with TEQUING exhibiting the highest value. No significant response to e-CO_2_ of plant leaf DM and SLA was observed (Table 2), although small but significant CO_2_ and cultivar effects were observed on plant green leaf area (P<0.01, data not presented).

Interestingly, a negative correlation was observed between local C source-sink ratio and the response to e-CO_2_ of total shoot DM (Fig. 4A, R^2^=0.67, P<0.05) and grain yield (Fig. 4B, R^2^=0.76, P<0.05). In both cases, a high local source-sink ratio was associated with a smaller positive response to e-CO_2_.

**Fig. 4:**
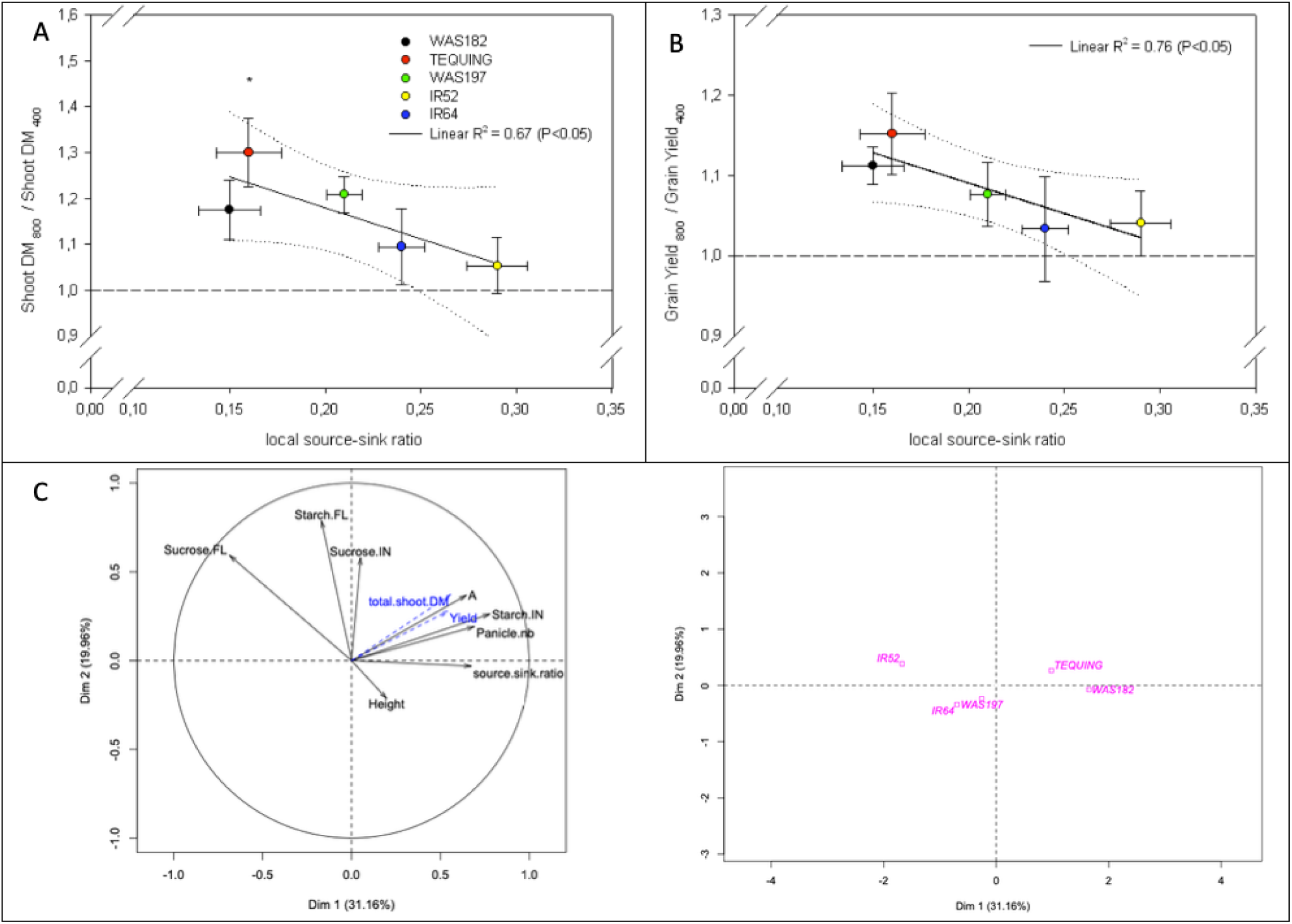
**(A): Relationship between total shoot biomass dry matter (DM) (stems + leaves) response to e-CO_2_ and local C source-sink ratio measured at ambient [CO_2_] in EXP2.** Average values +/- SE are presented. The average is made on 5 replicates, each replicate being computed as the ratio between shoot DM of a plant grown at e-CO_2_ (shootDM _800_) and the average (of 5 replicates) shoot DM of plants grown at ambient CO_2_ (shootDM _400_) **(B): Relationship between Grain Yield response to e-CO_2_ and local C source-sink ratio measured at ambient [CO_2_] in EXP2.** Average values +/- SE are presented. The average is made on 5 replicates, each replicate being computed as the ratio between grain yield of a plant grown at e-CO_2_ (Grain Yield 800) and the average (of 5 replicates) grain yield of plants grown at ambient CO_2_ (Grain Yield _400_) Stars indicate significant differences at *p* < 0.05 (Tukey HSD test) compared to ambient (dashed line). Each point represents the mean, in each CO_2_ treatment, of 5 values ± SE **(C): PCA using morphological** (Total shoot DM, grain Yield, Panicle number, local source-sink ratio, plant height), **biochemical variables** (Sucrose and Starch in the Flag leaf FL and the corresponding Internode IN), and **net photosynthetic rate** (*A*_*400*_) **for EXP2**. The centroid for each genotype is positioned (magenta squared) on the right part.

### Nonstructural carbohydrate (NSC) response to e-CO_2_ treatments

In both experiments, a strong cultivar effect (P<0.001) was observed on NSC contents (sucrose, hexose, starch) in both flag leaves and internodes. In EXP1 an additional [CO_2_] interaction effect was observed on flag leaf sucrose content. This was associated with a significant increase by +33% and +24% for IR52 and IR64, respectively, whereas LSS cultivars did not show any variation. Conversely, a [CO_2_] effect (Table 1, P<0.001) was observed on flag leaf starch content. This effect was positive in all cultivars. Regarding stem starch concentration, no significant effect of [CO_2_] was observed, although there was a trend of increased internode starch content under e-CO_2_ in LSS cultivars (+25% on average), whereas it was stable in the HSS cultivar.

In EXP2, a [CO_2_] effect was observed (Table 2, P<0.05) on leaf sucrose content, with a large increase by 40% and 17% in IR52 and IR64, respectively. With regards to stem sucrose and hexose contents, as in EXP1, no [CO_2_] or interaction effects were noticed. [CO_2_] generally increased leaf starch concentration (Table 2, P<0.01). Finally, as observed in EXP1, an increase in stem starch content by 24% on average was observed in LSS cultivars, whereas it numerically decreased by 32% in the HSS cultivar (not significant).

A principal component analysis (PCA) was performed (Fig. 4C) in order to relate the response to e-CO_2_ of shoot DM and grain yield to that of *A*_*400*_, morphological traits and biochemical traits in EXP2. The first dimension of the PCA (Dim 1) explained 31.2 % of the variation observed and was positively correlated to the response to CO_2_ of internode starch, panicle number, local source-sink ratio, *A*, total shoot DM and yield (r of 0.77, 0.69, 0.67, 0.64, 0.55, and 0.54 respectively). It was negatively correlated to the response of flag leaf sucrose (r = −0.68). Dim 2 explained 20.0 % of the variation and was correlated with flag leaf starch and sucrose content (r of 0.78 and 0.59, respectively) and internode sucrose content (r = 0.57). The genotypes were mainly discriminated by Dim1, with LSS and HSS cultivars well differentiated. Starch in internodes was the most discriminant biochemical variable, LSS genotypes exhibiting the greatest increase under e-CO_2_. Conversely, LSS showed the smallest increase of flag leaf sucrose under e-CO_2_.

## Discussion

A recent field study (Kikuchi *et al.*, 2017) suggested that rice genotypic differences in growth and yield responses to e-CO_2_ depend on the phenotypic plasticity of tillering and panicle size, giving plastic cultivars the capability of adjusting sink capacity to the increased assimilate source. Also, Fabre *et al.* (2019) demonstrated that the removal of the panicle sink causes strong reductions in rice photosynthesis particularly in the afternoon, and this inhibition was even more pronounced in e-CO_2_ treatments which enhanced the source-sink imbalance. Our results confirm that constitutive morphological differences, presumably affecting source-sink ratio in rice (our proxy parameter flag-leaf / panicle size ratio), affect the plant’s ability to efficiently use increased CO_2_ resources. We called this morphological proxy parameter a “local C source-sink ratio”.

### Local source-sink ratio: a driver of leaf photosynthetic capacity under e-CO_2_

Our study showed that the flag leaf photosynthesis (*A*_*400*_) of rice cultivars with low local C source-sink ratio (LSS) responded positively to e-CO_2_ whereas this was not the case for HSS genotypes (Fig. 1A). In addition, a strong negative correlation was observed across cultivars between the response of photosynthetic capacity to e-CO_2_ (*Amax*_*800*_/*Amax*_*400*_ of flag leaf) and the local C source-sink ratio (Fig. 2).

Variation in *Amax* for a given genotype, at a given level of light acclimation and abundant N resources, is primarily indicative of variation in *TPU* capacity, which in turn is the photosynthetic parameter linking the production and the consumption (or export) of fresh assimilates (Yang *et al.*, 2016; Fabre *et al.*, 2019). Sharkey (2016) stated that “When the highest data points of the *A*/*C*_i_ curve do not increase with CO_2_, *TPU* limitation is assumed”. In fact, a decline in φPSII should also occur at increasing *C*_i_ when *TPU* is limiting (Sharkey *et al.*, 1986; Long and Bernacchi, 2003). This condition was observed in IR52 and IR64 under e-CO_2_, but not in LSS cultivars (Fig. 3B). The low *Amax* response to e-CO_2_ of genotypes having high local C source-sink ratio might therefore indicate a lower *TPU* capacity in these cultivars, limiting *Amax*. Different levels of sink limitation as indicated by local C source-sink ratio would thus be reflected by differences in flag leaf *TPU* capacity.

Although high local source-sink ratio limited *A*_*400*_ and *Amax* under e-CO_2_ in HSS and intermediate cultivars, down regulation of photosynthesis was not necessarily directly caused by *TPU* limitation, as also reported by Fabre *et al.* (2019). A decrease in carboxylation efficiency (*V*_cmax_), stomatal conductance and other biochemical parameters may have participated (von Caemmerer and Farquhar, 1984; Shimono *et al.*, 2010; Zhu *et al.*, 2014). Co-adjustments of *V*_cmax_, *J*_max_ and *TPU* are common as these parameters vary together in strict stoichiometry (McClain and Sharkey, 2019). No significant reduction of leaf N or chlorophyll content was observed (Table 1). These factors are known to affect leaf photosynthesis under continuous e-CO_2_ conditions (Nakano *et al.*, 1995; Ainsworth and Long, 2005). Neither did we observe any K deficiency that potentially affects photosynthesis through stomatal closure (Wang *et al.*, 2013) and inhibited of assimilate export from leaves, causing assimilate accumulation in the leaf (Gerardeaux *et al.*, 2010).

A decline of photosynthesis under sink limitation, as observed here in HSS cultivars, has often been attributed to end-product accumulation in photosynthetic tissues (Paul and Pellny, 2003). Our data, in line with previous studies on rice (Shimono *et al.*, 2010; Fabre *et al.*, 2019), confirm a greater increase in leaf sucrose content in HSS than LSS cultivars (Table 1). The local excess of assimilates may negatively feedback on photosynthetic rate (Huber and Huber, 1992; Iglesias *et al.*, 2002; Li *et al.*, 2015; Yang *et al.*, 2016), possibly *via* leaf *TPU* capacity (Paul and Foyer, 2001; Fabre *et al.*, 2019).

Flag leaf starch content was also significantly increased under e-CO_2_, in line with previous findings under field conditions (Zhu *et al.*, 2016). However, the values observed (below 100 μg cm^-2^) were smaller than the empirical concentration (600 μg cm^-2^) reported to affect photosynthesis in rice (Weng and Chen, 1991). Thus, this parameter was probably not responsible for limited photosynthetic rates under e-CO_2_ here. The LSS cultivars also accumulated more starch in stem internodes under e-CO_2_. Stems are alternative or complementary sinks (Morita *et al.*, 2016) and may thus contribute to a more balanced source-sink ratio under e-CO_2_ at whole-plant scale. Indeed, the LSS cultivars had greater stem DM.

Increasing the photosynthetic capacity of the flag leaf has been reported to be an important target to improve rice yield potential particularly under e-CO_2_ (Chen *et al.*, 2007). According to our results, it is essential to simultaneously improve panicle C sink capacity to avoid photosynthetic efficiency losses caused by sink limitation.

### Impact on dry matter accumulation and grain yield

Efforts were made to design rice varieties more productive under long-term e-CO_2_, in particular by improving of photosynthesis (Raines, 2011). Dry matter production is stimulated by e-CO_2_ conditions (Shimono *et al.*, 2008, 2009; Ainsworth, 2008; Roy *et al.*, 2012) whereby the additional assimilates are accommodated by the plant structure in many species and cultivars by increased branching (Ziska *et al.*, 2004; Hasegawa *et al.*, 2013; Kirschbaum and Lambie, 2015). In the present study, we observed an increase of tillering in response to long-term e-CO_2_, particularly for LSS cultivars (Fig. S2). This is also in line with (Kikuchi *et al.*, 2017) who observed that rice genotypes with high tillering capacity respond strongly to e-CO_2_. Tillering in rice is known to be highly plastic and largely resource driven (Kumar *et al.*, 2016, 2017), and plant C supply-demand ratio is used to predict tillering in several crop models (Luquet *et al.*, 2006). The plasticity of the panicle sink is also related to branching of the rachis which generates new spikelet positions. It thus can be expected that at e-CO_2_, plant shoot apex activity in terms of cell division rates and branching will increase during the tillering phase (Seneweera and Conroy, 1997). In line with this, LSS genotypes exhibited a greater increase of tillering compared to HSS genotypes under long-term e-CO_2_. This result suggests that genotypes with the higher constitutive sink capacity also had greater sink plasticity. In fact, studies reported a close correlation between adaptation to CO_2_ enrichment and higher sink plasticity (Atwell *et al.*, 1999; Dahal *et al.*, 2014; Burnett *et al.*, 2016; Kikuchi et al., 2017). However, to our knowledge the genotypic covariation between sink strength and plasticity was never pointed out, in particular for traits involved in different phenological phases. The C source-sink limitation of crop growth under e-CO_2_ has been mainly studied during the reproductive phase (Peterhansel and Offermann, 2012; Lawlor and Paul, 2014; White *et al.*, 2016). This result may have implications for breeding for future climate conditions.

The local C source-sink ratio did not vary between EXP1 and EXP2 or between control and e-CO_2_ treatments (Fig. S1); Its component traits flag leaf area and panicle spikelet number both increased slightly under long-term CO_2_ exposure. Cultivars probably maintained this trait ratio at a genotypic level as suggested by (White *et al.*, 2016). The plasticity of flag leaf area and panicle spikelet number was greater in LSS than in HSS cultivars. Greater sink plasticity and sink potential in LSS were associated with higher starch accumulation in internodes (Table 2), similar to findings of (McKinley *et al.*, 2018). However, differences between LSS and HSS genotypes in sugar concentrations were not always significant and should be validated for a larger range of genotypes and/or CO_2_ conditions. Flag leaf hexose concentration was significantly increased by e-CO_2_ (Table 2); its positive role in sink strength and sink development was previously reported (Kumari and Asthir, 2016).

Long-term CO_2_ enrichment (EXP2) increased plant grain yield (Table 2, P<0.01) across cultivars, although the effect was not significant for any given cultivar. This commonly observed effect is related to increased number of fertile tillers and grains per panicle (Liu *et al.*, 2008; Nakano *et al.*, 2017). However, the significant dependency (P<0.05) of e-CO_2_-induced grain yield gain on local source-sink ratio, with LSS cultivars being more responsive than HSS, is new. Studies on wheat demonstrated a positive relationship between grain yield and photosynthesis of the flag leaf during grain filling (Reynolds *et al.*, 2000; Furbank and Parry, 2013). Our results suggest that grain yield gains through increased carbon assimilation require a concomitant increase of the panicle sink, and that rice varieties differ in this respect. Other authors have pointed out that flag leaf photosynthesis and sink capacity should be improved conjointly in rice (Shimono *et al.*, 2009; Hasegawa *et al.*, 2013, 2016; Ludewig and Sonnewald, 2016; Nakano *et al.*, 2017) and wheat (Reynolds *et al.*, 2017).

Rice grain filling relies on not only newly formed assimilates, but also on remobilized stem NSC reserves. Storage chiefly takes place until heading (Murchie *et al.*, 2009; Wang *et al.*, 2016), followed by remobilization (MacNeill *et al.*, 2017) which compensates for declining photosynthesis during late grain filling period (Yang and Zhang, 2006). Stem storage activity may resume when the grains are filled. Stems thus undergo a sink to source status transition (Cock and Yoshida, 1973; Chen and Wang, 2008). In the present study, there was a trend towards increased internode starch concentration under e-CO_2_ conditions at 15 days after heading for LSS cultivars, although not significant for any given cultivar. It thus seems that LSS cultivars mobilized stem reserves less exhaustively under e-CO_2_. The LSS cultivar TEQUING had a remarkably high level of non-mobilized stem reserves in both experiments. The present data do not allow interpreting the role of stem reserves in the response of LSS and HSS cultivars to e-CO_2_, and more research is needed on the regulation of reserve accumulation and remobilization variable source-sink relations.

Our study provided evidence that the genotypic local C source-sink ratio, as well as the plasticity of its component traits flag leaf and panicle size, is important for the ability of rice to translate enhanced CO_2_ resources into grain production. This information, in combination with previous findings on the importance of compensatory tillering ability (Kikuchi *et al.*, 2017), provides potential inroads for breeding for higher grain yield under the enhanced CO_2_ resources brought about by anthropogenic climate and atmosphere changes.

### Implications for crop modelling and phenotyping

Our data suggest that modelling leaf photosynthesis and its genotypic and environmental variability should consider source-sink feedbacks, particularly where e-CO_2_ conditions increase the C source-sink ratio. We observed that some simple morphological traits such as flag leaf size and panicle sink size, here used to calculate a proxy parameter for local source-sink ratio, affect photosynthetic and even grain yield response to e-CO_2_ under controlled conditions. These traits and their phenotypic plasticity, as well as the plasticity of tillering (Kikuchi *et al.*, 2017), can be simulated by some crop models, e.g. (Kumar *et al.*, 2016, 2017).

Feedback effects of sink limitation on photosynthesis may in part explain why inconsistencies are sometimes observed when simulating photosynthetic capacity (Lombardozzi *et al.*, 2017, 2018; Fatichi *et al.*, 2019). In general, even crop models simulating photosynthesis mechanistically, e.g., GECROS (Yin and van Laar, 2005; Yin and Struik, 2017) do not account for biochemical feedback inhibitions of photosynthesis, such as via *TPU* (Sharkey, 1985; Busch *et al.*, 2018). However, as our study indicated, the interactions between sources and sinks determining crop response to e-CO_2_ conditions may in part be predicted on the basis of simpler traits while avoiding excessive model complexity. For this purpose, the present, new results need confirmation for a larger segment of genetic diversity and for field conditions.

The phenotyping of larger populations for traits affecting crop e-CO_2_ response, both at the levels of morphology and photosynthesis, will serve two applied objectives, (1) the species-wide validation of the traits that would justify their inclusion in generic crop models; and (2) the study of the traits’ genetics for the purpose of breeding for cultivars that are more productive under future climatic scenarios (Wang *et al.*, 2016).

## Conclusion

This study provided new insights into the role local C source-sink balance, C sink strength and plasticity at different biological scales as rice plants respond to e-CO_2_. It demonstrated that morphological traits affecting plant source and sink capacity can drive the genotypic responses of photosynthesis at leaf level, and thereby influence plant production. It suggested that for the grain filling phase, the local C source-sink ratio is a potential target for selecting more CO_2_-responsive cultivars, pending validation for a broader genotypic spectrum and for field conditions. We thus call for broader phenotyping efforts for such traits and their effect on crop e-CO_2_ response, providing genetic information to breeders and a rationale for improving crop models to better predict crop responses to e-CO_2_.

## Acknowledgments

This work was financially supported by CIRAD, French Agricultural Research Centre for International Development; http://www.cirad.fr). We thank Audrey Dardou, Christian Chaine, Remy Michel for their valuable assistance in technical support, and the biochemical phenotyping platform from AGAP research unit of CIRAD.

## Supplementary data

**Fig. S1:** Relationship between local C source-sink ratio (flag leaf area / fertile spikelet number on the main stem) measured after a medium-term (15d from heading, EXP1) and a long-term (EXP2) CO_2_ enrichment at 800 μmol mol^-1^ of five rice cultivars. Each point represents the mean, in each CO_2_ treatment, of 5 replicates ± SE

**Fig. S2:** Dynamics of tillering of 5 rice cultivars grown under ambient (400 μmol mol^-1^, white filled color) and elevated CO_2_ (800 μmol mol^-1^, black filled color with red edge) during EXP2. Each point is an average of 5 replicates per genotype and treatment at a given date.

**Table S1:** Description of the 5 cultivars selected within the PRAY diversity panel for the present study: Values are mean of the two years (2013, 2014) in field experiments at CIAT (Columbia). Local C source-sink ratio computed as flag leaf area divided by fertile spike number on the main stem.

